# Ets1 and IL17RA cooperate to regulate autoimmune responses as well as skin immunity to Staphylococcus aureus

**DOI:** 10.1101/2023.06.19.545307

**Authors:** Michael Battaglia, Alex C. Sunshine, Wei Luo, Richard Jin, Alifa Stith, Matt Lindemann, Lloyd S. Miller, Satrajit Sinha, Elizabeth Wohlfert, Lee Ann Garrett-Sinha

**Affiliations:** Department of Biochemistry, State University of New York at Buffalo, Buffalo, NY 14203, USA; Department of Microbiology and Immunology, State University of New York at Buffalo, Buffalo, NY 14203, USA; AESKU Diagnostics, Buffalo, NY 14203, USA; Department of Department of Dermatology, Johns Hopkins University School of Medicine, Baltimore, MD 21231, USA.

**Author notes:** Co-first authors (Michael Battaglia, Alex C. Sunshine). Corresponding author Lee Ann Garrett-Sinha Department of Biochemistry State University of New York 955 Main St, Rom 5124 Buffalo, NY 14203.

**Keywords:** autoimmunity, immunodeficiency, *Staphylococcus aureus*, autoantibodies, Ets1, IL17RA, dendritic epidermal T cell

## Abstract

Ets1 is a lymphoid-enriched transcription factor that regulates B and T cell functions in development and disease. Mice that lack Ets1 (Ets1 KO) develop spontaneous autoimmune disease with high levels of autoantibodies. Naïve CD4+ T cells isolated from Ets1 KO mice differentiate more readily to Th17 cells that secrete IL-17, a cytokine implicated in autoimmune disease pathogenesis. To determine if increased IL-17 production contributes to the development of autoimmunity in Ets1 KO mice, we crossed Ets1 KO mice to mice lacking the IL-17 receptor A subunit (IL17RA KO) to generate double knock out (DKO) mice. We found that the absence of IL17RA signaling did not prevent or ameliorate the autoimmune phenotype of Ets1 KO mice, but rather that DKO animals exhibited worse symptoms with striking increases in activated B cells and secreted autoantibodies. This was correlated with a prominent increase in the numbers of T follicular helper (Tfh) cells. In addition to the autoimmune phenotype, DKO mice also showed signs of immunodeficiency and developed spontaneous skin lesions colonized by *Staphylococcus xylosus*. When DKO mice were experimentally infected with *S. aureus* they were unable to clear the bacteria, suggesting a general immunodeficiency to Staphylococcal species. γδ T cells are important for control of skin Staphylococcal infections. We found that mice lacking Ets1 have a complete deficiency of the γδ T cell subset dendritic epidermal T cells (DETC), which are involved in skin wound healing responses. To determine if loss of DETC might promote susceptibility to Staph infection, we depleted DETC from IL17RA KO mice and found that the combined loss of DETC and IL-17 signaling leads to a failure to clear the infection. Our studies suggest that defects in wound healing, such as that caused by loss of DETC, can cooperate with impaired IL-17 responses to lead to increased susceptibility to skin Staph infections.

## Introduction

The Ets1 transcription factor is highly expressed in B and T cells and regulates their differentiation (1). Mice lacking Ets1 (Ets1 KO mice) develop autoimmune disease, with similarities to human lupus (2). Indeed, genome-wide association studies (GWAS) have associated single nucleotide polymorphism (SNPs) in the human *ETS1* gene locus with an increased susceptibility to SLE (3–5). The levels of Ets1 mRNA are reduced in PBMCs of lupus patients and are inversely correlated with the serum titers of autoantibodies against dsDNA (4, 6, 7). SNPs in the *ETS1* locus are also associated with susceptibility to rheumatoid arthritis, psoriasis and ankylosing spondylitis, as well as multiple other autoimmune and inflammatory diseases (8–13).

The autoimmune phenotype of Ets1 KO mice has been attributed to changes in both the B and T cell compartments. Indeed, Ets1 KO mice have increased percentages of activated B cells, class-switched B cells and antibody-secreting plasma cells (2, 14, 15). This is coupled with increased levels of serum IgM, IgG1 and IgE (16, 17). Serum of Ets1 KO mice also has high titers of autoantibodies to double-stranded DNA and other autoantigens (2, 17). *In vitro*, Ets1 KO B cells have been shown to differentiate more readily into plasma cells when exposed to Toll-like receptor stimulation (2, 18, 19). The increase in plasma cells *in vivo* and production of autoantibodies has a B cell-intrinsic component, as shown by generation of mixed bone marrow chimeras and B cell specific knockout of Ets1 (18, 20).

In the T cell compartment, Ets1 KO mice have a number of important alterations, including an increase in the number of T cells with a memory phenotype (17, 21), increased differentiation into T follicular helper type 2 (Tfh2) cells that secrete high levels of IL-4 (22) and enhanced generation of Th17 cells (23). Furthermore, SLE patients carrying Ets1 risk alleles tend to have higher serum IL-17 levels than patients lacking these risk alleles (24). Ets1 KO mice have also been reported to have fewer Foxp3+ CD25+ T-regulatory cells in the spleen and thymus and the Treg cells that do develop have impaired suppressive activity (17). This is likely due to a role for Ets1 in binding to regulatory elements in the *Foxp3* gene to promote transcription (17, 25). Together, these alterations in T cell differentiation contribute to progression of autoimmune disease in Ets1 KO mice, as evidenced by knockout of Ets1 specifically in T cells, which results in autoimmune disease (22).

Th17 cells and IL-17 have both been shown to be increased in multiple autoimmune diseases and to play roles in driving inflammatory pathogenesis. In SLE, increased levels of IL-17 have been positively correlated with disease severity and negatively correlated with response to immunosuppressive treatment (26). In addition to its described role in autoimmune diseases, IL-17 is also important in immune responses against pathogens such as *Staphylococcus aureus* (*S. aureus*) (27) and *Candida albicans* (*C. albicans*) (28). The protective role of IL-17 against certain pathogens is ascribed to its ability to induce production of antimicrobial peptides that kill bacteria and chemokines involved in recruiting neutrophils (29–31). To test the role of IL-17 in promoting autoimmune disease in Ets1 KO mice, we crossed Ets1 KO mice to IL17RA KO mice to generate double knock out (DKO) mice. The resulting mice can produce IL-17, but are unable to respond to it. Given the aforementioned role of IL-17, we anticipated that autoimmune disease in these mice might be less severe than in Ets1 KO mice. However, to our surprise, DKO mice displayed worse autoimmune disease than control mice and this was coupled with increased numbers of Tfh cells, germinal center B cells, class-switched B cells, memory B cells and plasma cells. Furthermore, they demonstrated a high susceptibility to Staphylococcal infections of the skin. γδ T cells are known to be crucial for anti-Staphylococcal skin immune responses. We found that mice lacking Ets1 lack the dendritic epidermal T cell (DETC) subset of γδ T cells. To probe the mechanistic role of DETC in Staphylococcal infection, we depleted DETC from IL17RA KO mice, which led to enhanced susceptibility to skin Staphylococcal infections. Thus DETC, whose development is dependent on Ets1, cooperate with IL-17 signaling to regulate skin immune responses to Staphylococcal infection. The persistent skin colonization with high levels of Staph bacteria likely induces increased immune cell activation and may contribute to the development of increased auto-immune responses in DKO mice as compared to Ets1 KO mice.

## Methods

### Mice

The following mouse strains were used in this report: Ets1^-/-^ (Ets1 KO) (2), IL17RA^-/-^ (IL17RA KO) (32) and Ets1^-/-^IL17RA^-/-^ (DKO mice) as well as wild-type littermate controls that were obtained by breeding heterozygotes of the above strains. IL17RA KO mice used in this study were obtained from Amgen, Inc (31). Since the loss of Ets1 results in perinatal lethality on an inbred C57BL/6 background (33), all mice were maintained on a mixed C57BL/6 x 129Sv genetic background. Blood was obtained under anesthesia using either retro-orbital bleed or cardiac puncture. Animals used for experiments were euthanized with CO_2_ induction followed by cervical dislocation. Mice were housed under specific pathogen free (SPF) conditions for the duration of the studies. All studies were approved by the University at Buffalo IACUC committee.

### Flow cytometry

Spleen and lymph nodes were isolated from wild-type, Ets1 KO, IL17RA KO and DKO mice and single cell suspensions were prepared. Cells were incubated with Ghost Dye Violet 510 (Tonbo Biosciences, San Diego, CA) to stain dead cells. Cells were subsequently stained with surface antibodies for B and T cell marker proteins and by intracellular staining for key transcription factors. Fluorescent signals were collected with an LSRII or Fortessa flow cytometers. Antibodies for flow cytometry were obtained from BD Biosciences, BioLegend, eBioscience, Miltenyi Biotec or R&D Systems. The following antibodies were used in flow analysis of B cell subsets: B220 (clone RA3-6B2), CD21 (clone 7G6 or REA800), CD23 (clone B3B4), CD73 (clone eBioTY/11.8), CD80 (clone 16-10A1), CD138 (clone 281-2), FAS (clone Jo2), IgG1 (clone RMG1-1), PD-L2 (clone TY25) and peanut agglutinin (PNA, biotinylated from Vector Labs, Burlingame, CA). The following antibodies were used in flow analysis of T cell subsets: CD4 (clone GK1.5), CD8b (clone eBioH35-17.2), PD1 (clone 29F.1A12) and CXCR5 (clone L138D7). These antibodies were used for intracellular staining of key transcription factors regulating T helper cell subset differentiation: Tbet (clone 4B10), Foxp3 (clone FJK-16s), RORγT (clone B2D), GATA3 (clone L50-823) and BCL6 (Clone 7D1). For isolation of skin associated lymphocytes, depilated whole skin was harvested and floated on EDTA-free trypsin (ThermoFisher) overnight at 4°C. The epidermis and dermis were then separated. The epidermal sheets were digested further in trypsin supplemented with .01% DNAse I (Millipore Sigma). The dermis was digested with 0.25 mg/ml Liberase (Millipore Sigma) and 1 mg/ml DNAse I (Millipore Sigma). The resulting cell suspensions were enriched for lymphocytes using Lymphoprep (Stemcell Technologies). Following enrichment, cells from the dermis and epidermis were combined and cultured overnight in the presence of 10 U/ml IL-2 (Biolegend). The following antibodies were used in flow analysis of skin associated γδ T cell populations: CD4 (clone GK1.5), CD8b (clone eBioH35-17.2), TCRγδ (clone GL3), Vγ5 (clone 536) and CCR6 (clone 29-2L17).

### *In vitro* differentiation of Tfh cells

Spleen and lymph nodes from mice were harvested, mechanically disrupted, and passed through a 70μm filter. Naïve CD4^+^ T cells from the sample were purified by magnetic bead separation using the CD4^+^CD62L^+^ isolation kit (Miltenyi Biotec). Cells were plated in 96-well plates coated with anti-CD3 antibody (5μg/ml, BD, clone 145-2C11) at a density of 3 x 10^6^ cells/ml. Cells were plated in Tfh conditioning media (RPMI complete media with 10% fetal bovine serum, 10μg/ml anti-IL4 (BD Biosciences, clone 11B11), 10μg/ml anti-IFNγ (BD Biosciences, clone XMG1.2), 10μg/ml anti-TGFβ (R&D Systems, clone 1D11), 30ng/ml IL6 (Shenandoah Biotechnology), 50ng/ml IL21 (Shenandoah Biotechnology) in the presence of soluble anti-CD28 (2μg/ml, BD, clone 37.51). Polarized cells were harvested for Tfh phenotyping by flow cytometric analysis 4 days after plating.

For surface phenotype and transcription factor analysis, harvested cells were stained Live/Dead Fixable Aqua dead cell stain (Thermo Scientific Fisher) to exclude dead cells. Cells were subsequently stained for surface markers (CD4, TCRβ, PD1, CXCR5) and then fixed and permeabilized using the Intracellular Fixation and Permeabilization Buffer Set (eBioscience). Fixed cells were stained with antibody to BLC6 (Clone 7D1) or isotype control.

### ELISA

ELISA to detect total serum IgM and IgG and autoantibodies was performed as previously described (2). Serum IgE levels were detected using the BioLegend Mouse IgE ELISA Max Deluxe Set). Maxisorp 96 ELISA well plates were coated overnight with 10μg/ml of antigens and ELISA was performed as previously described (2).

### ELISPOT

Single cell suspensions were prepared from spleen and lymph nodes and plated on ELISPOT plates to detect IgM- and IgG-secreting cells as previously described (2, 15). Spots were counted with an automated counter and the numbers of antibody-secreting cells per million total cells plated was calculated.

### Immunostaining of Hep2 cells

Hep2 hepatoma cells on glass slides were incubated with 1:40 dilutions of mouse serum and subsequently with FITC-conjugated anti-mouse IgG. Staining was performed and images were captured with a HELmed Integrated Optical System (HELIOS, AESKU Diagnostics).

### RNA-sequencing and analysis

Skin samples harvested from DKO and control mice were processed in TRIzol (Thermo Fisher) and bulk RNA was then purified. Purified RNA was then analyzed by QuBit/Quant-IT and Fragment Analyzer (Agilent) for quality control. cDNA library preparation was completed using TrueSeq RNA sample preparation kit (Illumina) followed by 50-bp single-end sequencing on an Illumina HiSeq 2500. Analysis of the raw reads using FASTQC v0.11.9 application was used as a quality control metric following sequencing.

The raw reads were then mapped to the GRCm38 (the mm10 reference mouse genome) using TopHat2 (34). The aligned reads were then quantified using featureCounts v1.5.3 to generate a raw counts matrix. The raw counts matrix was then processed in R to generate transcripts per million normalized expression values as previously described in Wagner et al (35). Differential gene expression analysis was conducted comparing knockout samples to wild-type controls using DEseq2 v1.24.0 with genes being identified as statistically significantly differentially expressed with a log2fold change ≥1 and a FDR value of ≤0.1.

Upregulated and downregulated DEGs identified by the above analysis were then subjected to GO term enrichment analysis. DEGs were input into the online tool g;Profiler (36) to assess for GO terms significantly enriched among the upregulated and downregulated genes.

### Analysis of susceptibility to *Staphylococcus* infection

To determine if DKO have spontaneous colonization of the skin with Staphyloccal bacteria, we swabbed the ventral side of the neck and upper thorax and plated swabbed bacteria on mannitol-salt-agar (MSA) plates that specifically promote growth of *Staphylococcal* species, while suppressing the growth of other bacteria. Plates were incubated overnight at 37°C and the colonies were subsequently counted. Genomic sequencing was used to confirm the species of Staphylococcal bacteria. To determine if DKO mice are susceptible to exogenous *S. aureus* infection, a methicillin-resistant *Staphylococcus aureus* (MRSA) USA300 strain NRS-384 carrying a bioluminescent marker (the lux operon from *Photorhabdus luminescens*) was obtained from Dr. Roger Plaut at the Food and Drug Administration (FDA) (37). Bacteria were grown in tryptic soy agar to log phase, harvested and washed with PBS and then resuspended at 200 million cells/ml. Mice to be infected were anesthetized followed by shaving the back skin and making three small topical cuts in the skin. Approximately 10 microliters of resuspended bioluminescent *S. aureus* was introduced into the cuts. Mice were given buprenorphine to control pain. Infected mice were imaged on days, 1, 3, 7, 14 and 21 post-infection using an IVIS imager that detects the bioluminescent signal. Luminescence values were normalized to the signal at day one to control for differences in cut size and depth between animals.

### Depletion of DETC

WT and IL17RA KO animals were injected with 100μg of Ultra-LEAF Purified Anti-mouse TCR Vγ5 antibody (Clone 536) or Ultra-LEAF Purified Syrian Hamster IgG Isotype control antibody (Clone SHG-1). For validation of depletion, epidermal sheets were harvested from ear leaflets as previously described (Jameson et al 2004). Epidermal sheets were stained with FITC-anti-mouse TCRγδ (Clone GL3) and TOPRO-3 Iodide (ThermoFisher). For determination of DETC contribution to skin *S. aureus* infection immune responses, WT and IL17RA KO animals were injected with antibody as above and infected two days post-injection as described above. The burden of *S. aureus* was measured using an IVIS imager on days 1, 2, 3, 4, 5, 7, 10, 14, 21, 24, 28, 32, and 35 days post infection with signal normalized to day 1 in order to control for cut size and depth variation.

### Statistics

One way ANOVAs or non-parametric Kruskal-Wallis tests were performed followed by Tukey’s or Dunn’s multiple comparison test respectively, except for the autoantibody ELISAs and IVIS imaging, where 2 way ANOVAs were used followed by a Bonferoni post-test. In all cases, error bars are standard error of the mean (SEM) and p<0.05 was considered significant.

## Results

### Loss of IL-17 signaling promotes autoimmune disease in Ets1 knockout mice

In order to test the role of IL-17 in the autoimmune phenotype of Ets1 deficient mice, we crossed Ets1 KO mice to mice lacking the IL-17 receptor A subunit (IL17RA) to generate double knockout mice (DKO). We analyzed DKO and control wild-type and single knockout mice at 3-6 months of age. While we anticipated that loss of IL-17 signaling might result in reduced immune cell activation and autoimmune phenotypes, we instead found that DKO mice had a stronger phenotype than Ets1 KO mice. This was shown by greatly enlarged peripheral lymph nodes (Figure 1A-B) and more modestly enlarged spleens in the DKO mice as compared to controls (Table I). ELISPOT assays showed that there was a dramatic increase in antibody-secreting cells in DKO mice compared to Ets1 KO mice, especially IgG-secreting plasma cells in the lymph node (Figure 1C-D). Levels of serum IgM and IgE are higher in Ets1 KO than wild-type controls, whereas total serum IgG titers are not significantly increased (16, 17). Serum from DKO mice showed a similar increase in IgM as that found in Ets1 KO serum, but had a much stronger increase in serum IgG and IgE (Figure 2A). On average, the serum IgE levels were >400 fold higher in DKO as compared to WT mice and ∼5 fold higher in DKO as compared to Ets1 KO.

**Figure 1:**
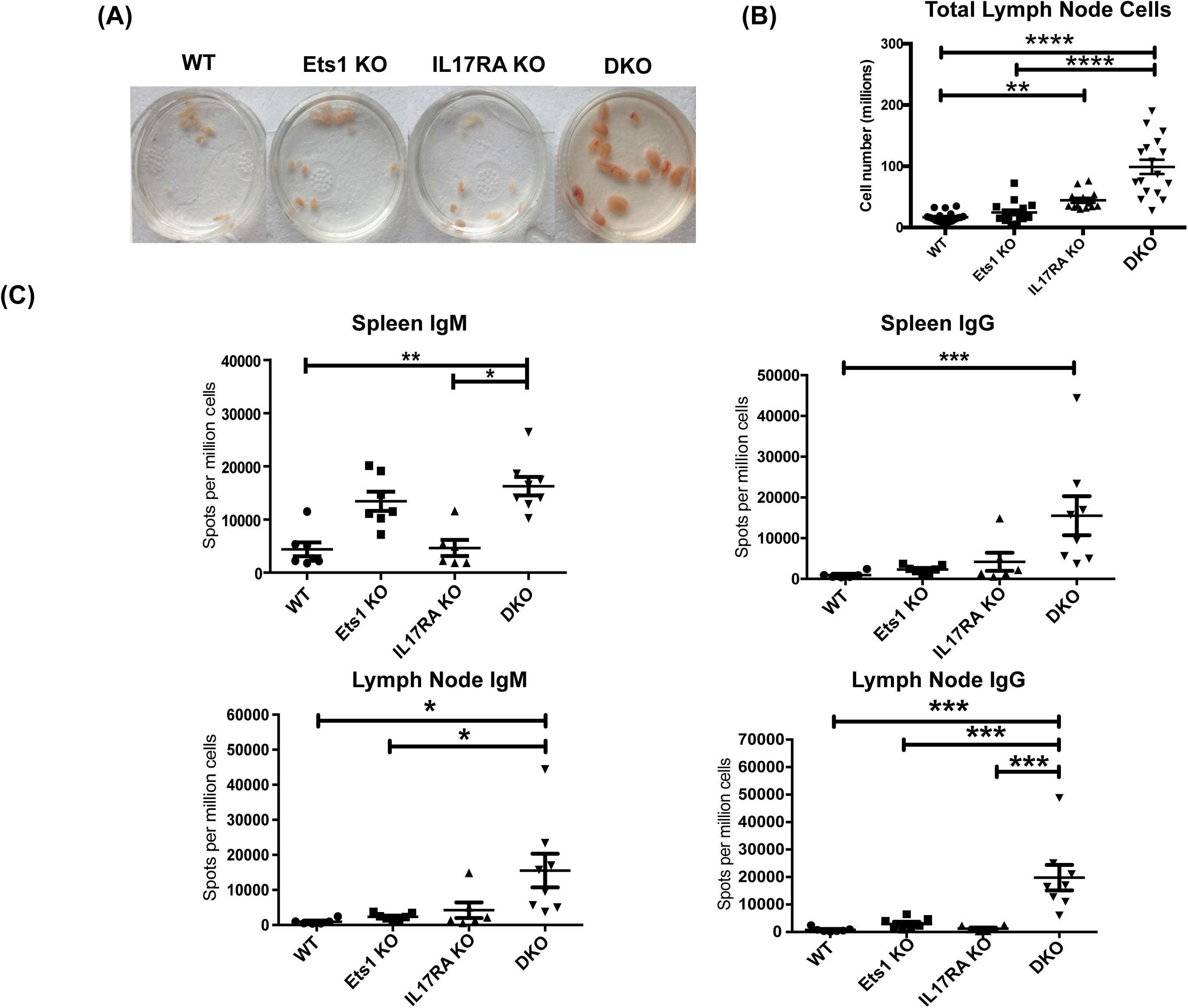
DKO mice have enlarged lymph nodes and increased plasma cells and serum antibodies. (A) Peripheral lymph nodes harvested from wild-type, Ets1 KO, IL17RA KO and DKO mice. (B) Total lymph node cells in mice of the indicated genotypes (n=13-19 mice per genotype). (C) ELISPOT quantification of IgM- and IgG-secreting plasma cells in spleen and lymph node of mice of the indicated genotypes (n=6-8 mice per group). *p< 0.05, **p<0.01, ***p<0.001, ****p<0.0001.

**Table I.** – Absolute Numbers of B and T Cell Populations in Spleen and Lymph Node

|  | Wild-type | Ets1<sup>-/-</sup> | IL17RA<sup>-/-</sup> | DKO |
| --- | --- | --- | --- | --- |
| Number of Splenocytes (millions) | 50.0 ± 4.9 | 83.3 ± 12.7 | 40.9 ± 3.6 | *** 111.4 ± 11.6 |
| Number of Lymphocytes (millions) | 16.6 ± 1.9 | 24.7 ± 4.5 | ** 44.3 ± 4.2 | **** 99.0 ± 11.7 |
| Number of B220<sup>+</sup> B cells (Spleen) | 25.08 ± 3.3 | 41.7 ± 8.0 | 17.34 ± 2.3 | ** 59.1 ± 5.9 |
| Number of B220<sup>+</sup> B cells (Lymph Nodes) | 3.45 ± 0.9 | 10.49 ± 3.37 | 7.47 ± 2.27 | **** 47.56 ± 7.34 |
| Percent of B220<sup>+</sup> B cells (Spleen) | 42.64 ± 3.12 | 46.42 ± 3.35 | 42.16 ± 3.62 | 50.6 ± 2.07 |
| Percent of B220<sup>+</sup> B cells (Lymph Nodes) | 17.36 ± 2.05 | *34.58 ± 4.09 | 14.88 ± 3.12 | *** 43.64 ± 2.49 |
| Number of CD4<sup>+</sup> T cells (Spleen) | 11.8 ± 1.43 | 19.42 ± 3.01 | 8.49 ± 0.99 | * 20.08 ± 2.33 |
| Number of CD4<sup>+</sup> T cells (Lymph Nodes) | 6.31 ± 0.82 | 7.97 ± 1.30 | *** 18.27 ± 2.11 | **** 21.17 ± 2.40 |
| Percent of CD4<sup>+</sup> T cells (Spleen) | 22.88 ± 2.65 | 23.92 ± 1.33 | 22.23 ± 2.01 | 18.85 ± 1.08 |
| Percent of CD4<sup>+</sup> T cells (Lymph Nodes) | 39.11 ± 3.03 | 36.38 ± 3.56 | 41.93 ± 3.21 | *** 22.91 ± 1.40 |

\* p<0.05, \*\*p<0.01, \*\*\*p<0.001, \*\*\*\*p<0.0001 (when compared to wild-type controls)

**Figure 2:**
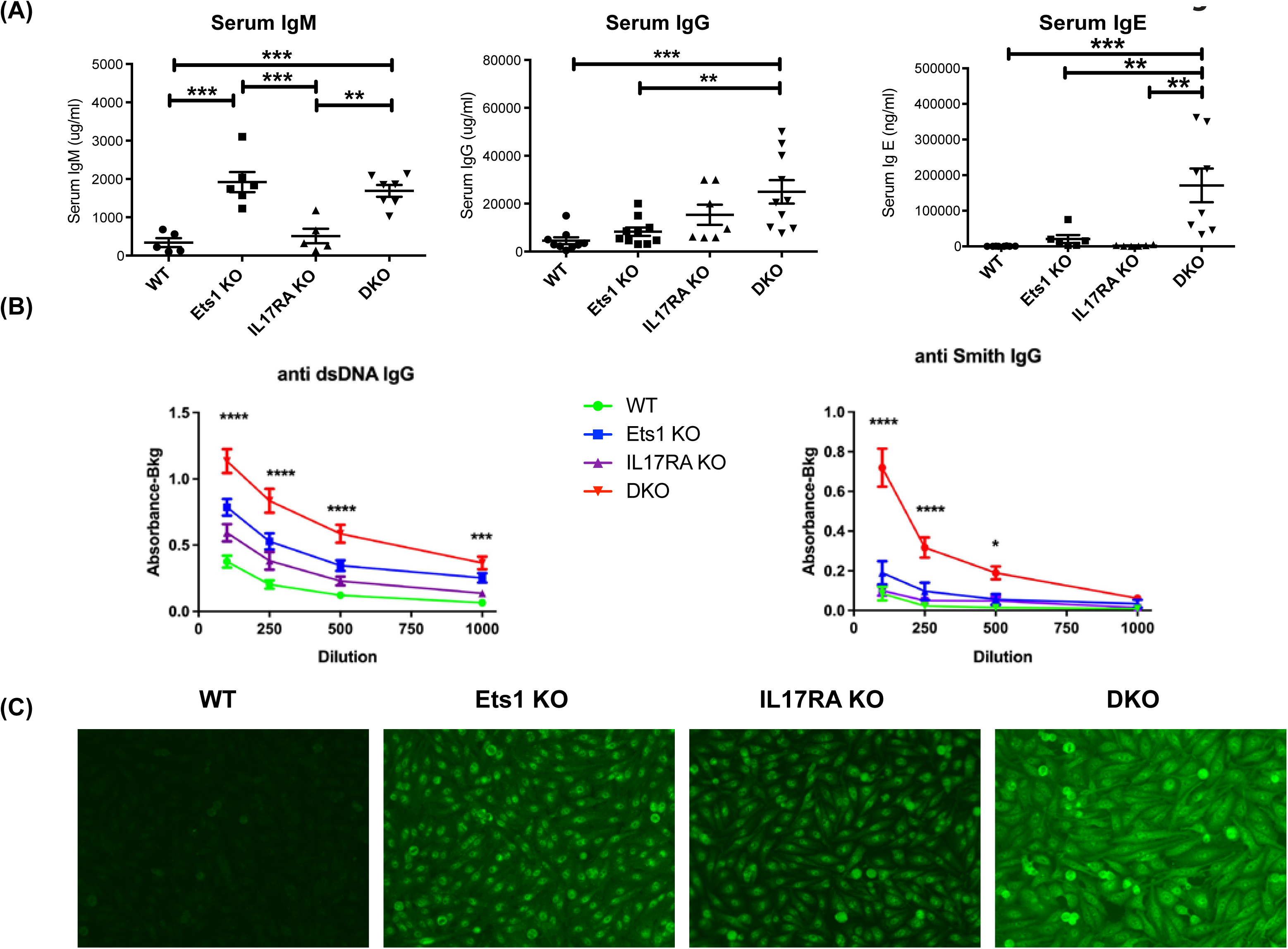
DKO mice have increased titers of autoantibodies. (A) ELISA quantification of total serum IgM, IgG and IgE of mice of the indicated genotypes (n=5-7 mice per group for IgM, n=7-10 mice per group for IgG, n=6-8 mice per group for IgE). (B) IgG autoantibodies to dsDNA and Smith antigen (Sm) in serum from wild-type (WT, n=15), Ets1 KO (n=12-13), IL17RA KO (n=9-10) and DKO mice (n=16-18). Asterisks represent statistical significance when comparing WT to DKO at each serum dilution. (C) A representative image of Hep2 cells stained with serum from mice of the indicated genotypes and with a FITC-conjugated anti-mouse IgG secondary antibody. *p< 0.05, **p<0.01, ***p<0.001, ****p<0.0001.

DKO mice had higher titers of autoantibodies against double-stranded DNA (dsDNA) and Smith antigen (Sm) than Ets1 KO mice (Figure 2B). Hep2 cell staining demonstrated that Ets1 KO and DKO mice produce anti-nuclear and anti-cytoplasmic IgG autoantibodies (Figure 2C). Unexpectedly, we also found that IL17RA KO mice produce autoantibodies as well. The intensity of Hep2 staining was in general stronger using serum from DKO mice than Ets1 KO or IL17RA KO, indicating increased levels of autoantibodies in DKO mice.

### DKO mice have more activated B and T cells than Ets1 KO mice

To further explore the phenotype of DKO mice, we analyzed B and T cell differentiation status using flow cytometry. DKO mice have increased total B cell numbers in the spleen and lymph nodes compared to controls (Figure 3A and Table I). Similarly, DKO mice showed a striking increase in the percentages and numbers of plasma cells and IgG1 class-switched B cells compared to Ets1 KO mice (Figure 3B-C and Supplemental Figures 1-2). Like Ets1 KO mice, DKO showed a strong reduction of marginal zone B cells in the spleen (Figure 3D). We found that there was an increase in the numbers of germinal center B cells (B220+PNA+Fas+) in the lymph nodes and an increase in the numbers of B cells with a memory phenotype (B220+CD80+PDL2+) in both the spleen and the lymph nodes of DKO mice as compared to controls (Figure 3E-F and Supplemental Figures 3-4). These results indicate that DKO mice have a greatly expanded germinal center response and an overall high level of B cell activation.

**Figure 3:**
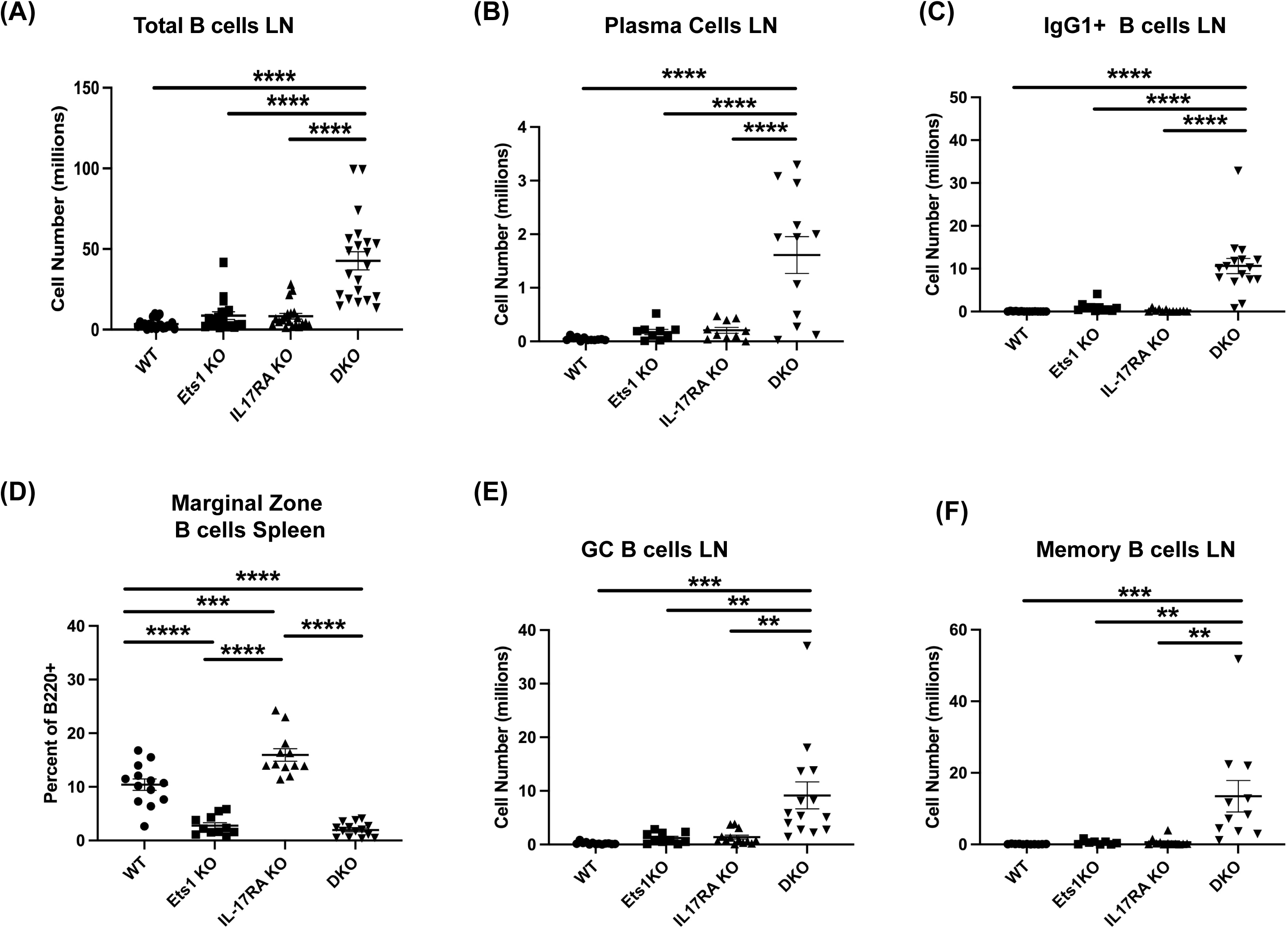
DKO mice have more activated and class-switched B cells. (A) Total B220+ B cells in lymph nodes of mice of the indicated genotypes (n=18-21 mice per genotype). (B) Quantification of the numbers of plasma cells (B220-low CD138+) (n=9-12 mice per genotype) and (C) IgG1+ B cells (B220+IgG1+) (n=11-16 mice per genotype) in lymph nodes. (D) Percent of CD21^hi^ CD23^low^ marginal zone B cells among total B220+ B cells in spleen (n=11-15 mice per genotype). (E) Quantification of germinal center B cells (B220+Fas+PNA+) (n=10-14 mice per genotype) and (F) memory phenotype B cells (B220+CD80+PDL2+) (n=8-11 mice per genotype) in the lymph nodes of mice of the indicated genotypes. * **p<0.01, ***p<0.001, ****p<0.0001.

**Figure 4:**
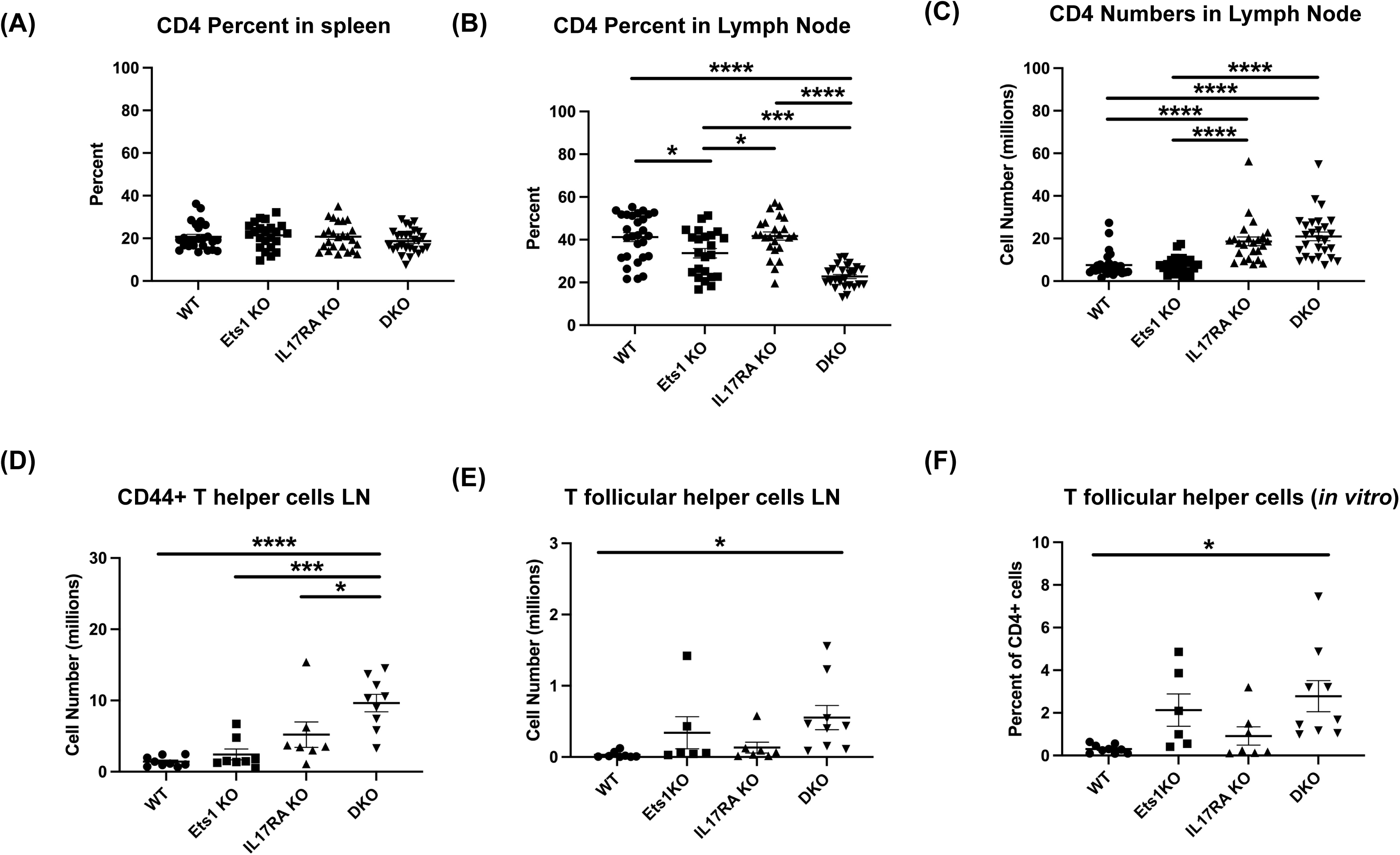
DKO mice have more activated T cells and increased numbers of Tfh. (A-B) Percent of CD4+ T cells in spleen and lymph nodes of mice (n=22-27 mice per genotype). (C) Total numbers of CD4+ T cells in the lymph nodes of mice (n=22-27 per genotype). (D) Quantification of the numbers of CD4+ T cells with an activated/memory phenotype in lymph nodes (CD4+CD44+) (n=7-9 mice per genotype). (E) Quantification of CD4+ T cells with a Tfh phenotype (PD1+CXCR5+) in lymph nodes (n=6-9 mice per genotype). (F) Quantification of the percentage of Tfh cells (PD1+CXCR5+ICOS+BCL6+) among total live CD4+ cells in *in vitro* cultures (n=6-9 mice per genotype). *p< 0.05, ***p<0.001, ****p<0.0001.

In the T cell compartment, we found that while the overall percentage of CD4+ T cells in the spleen was similar in all genotypes (Figure 4A), the percentage of CD4+ T cells in the lymph nodes of DKO mice was reduced (Figure 4B and Table I). However, because both IL17RA KO and DKO lymph nodes are larger than those of control mice, the total number of CD4+ T cells was in fact higher in IL17RA KO and DKO mice than in wild-type or Ets1 KO controls (Figure 3C and Table I). Many CD4+ T cells in the lymph nodes of IL17RA KO and especially DKO mice expressed CD44, a marker of an activated or memory phenotype (Figure 4D). Furthermore, in the CD4+ T cell subset, there was an increased number of cells with a T follicular helper (Tfh) phenotype (CD4+ PD1-hi CXCR5-hi BCL6+) in Ets1 KO and DKO spleen and lymph nodes (Figure 4E and Supplemental Figure 5). To further examine the propensity of DKO T cells to become Tfh, we isolated naïve CD4+ T cells from the spleens and lymph nodes of DKO and control mice and stimulated them *in vitro* under conditions that promote Tfh differentiation. CD4+ T cells from Ets1 KO and DKO mice showed increased differentiation to PD1+ CXCR5+ ICOS+ BCL6+ Tfh cells (Figure 4F and Supplemental Figure 6).

We also examined expression of Foxp3 in CD4+ T cells to assess the numbers of regulatory T cells (Tregs). A previous report had indicated that there was a 3-4 fold reduction in the percentages of Foxp3+ Tregs in the spleens of Ets1 KO mice (17). In our studies, we found that the number of Foxp3+ Tregs was similar in the spleens and lymph nodes of wild-type, Ets1 KO and IL17RA KO mice and was elevated in DKO mice (Supplementary Figure S7A-B). However, similar to the previous study (17), we did find that the intensity of Foxp3 staining was lower in CD4+ T cells from mice lacking Ets1, especially in the lymph node (Supplementary Figure S7C). We also analyzed staining of Tbet, GATA3 and Rorγt in spleen and lymph node cells to determine if there was spontaneous differentiation of CD4+ T cells to Th1, Th2 or Th17 fates. The total numbers of CD4+ T cells that stained with GATA3 and RORγt antibodies was elevated in DKO mice (Supplementary Figure S7C-D).

### DKO mice have increased susceptibility to Staphylococcal skin infections

We noted that by six months of age, most DKO mice began to develop skin dermatitis and lesions, often on the ventral surface of the neck (Figure 5A). Such lesions were not found on Ets1 KO mice and were found at a lower rate and lesser intensity in IL17RA KO mice. RNA-sequencing experiments using RNA isolated from the skin of DKO and control mice showed upregulation of many pathways associated with an inflammatory immune response, including the response to *Staphylococcus aureus* infection (Supplemental Figure 8A). Indeed, many cytokines and chemokines involved in the skin immune response were highly over-expressed in the skin of DKO mice as compared to control mice (Supplemental Figure 8B).

**Figure 5:**
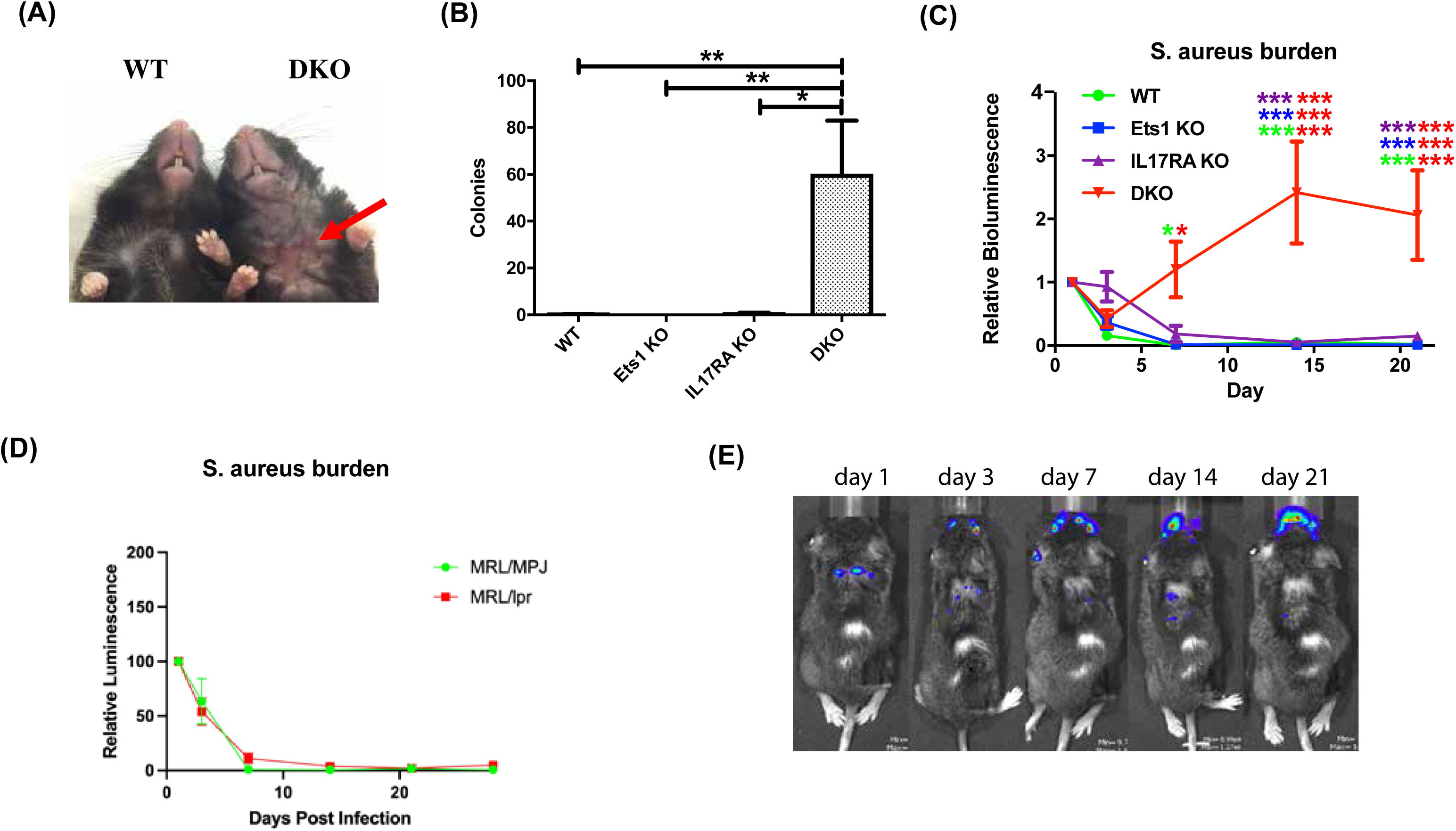
DKO mice are susceptible to skin infections with *S. aureus*. (A) Representative skin lesion on the ventral side of the neck of a DKO mouse (red arrow indicates lesion) compared to an unaffected wild-type (WT) control. (B) Quantification of the recovery of *S. xylosus* colonies from swabbing the ventral neck skin of mice of the indicated genotypes (n=4-7 per genotype). (C) Quantification of IVIS imaging of four-month-old DKO and control mice infected with bioluminescent *S. aureus*. Mice were imaged on days 1, 3, 7, 14 and 21 post-infection and signals were normalized to those on day 1. Asterisks are color coded for the strains compared (using the colors in the legend) and the number of asterisks of any one color above a particular point on the curve represents the significance of the difference at that time point and the two colors on each line of asterisks represent the two genotypes compared (n=5-7 per genotype). (D) Quantification of IVIS imaging of 4-month-old autoimmune MRL/lpr mice and non-autoimmune control MRL/MpJ mice. N=6 MRL/MpJ and 13 MRL/lpr mice. (E) Representative IVIS images of a DKO mouse showing shifting of the bioluminescent signal from the back, where the infection was initiated, to the face. *p< 0.05, ***p<0.001.

Mice lacking IL17RA are known to have a defect in clearing *Staphylococcus aureus* from the skin (27). Given this observation and the results of RNA-sequencing, we sought to determine whether Staphylococcal bacteria could be recovered from the skin of DKO. Swabbing the ventral neck of mice, resulted in the recovery of high levels of Staphylococcal bacteria from the skins of 100% of DKO mice tested (whether or not they showed visible lesions), while Staph was cultured from only two of six IL17RA KO mice tested using this technique. The level of Staph bacteria recovered from IL17RA KO mice was also lower than that found in DKO mice (Figure 5B). Genomic sequencing showed that recovered bacteria were *Staphylococcus xylosus* (Supplemental Figure 8C), a Staph species prominent on mouse skin (38–40) and a frequent cause of skin infections in mice (41–44).

To determine whether DKO mice have an impaired immune response to Staphylococcal bacteria, we experimentally infected cuts in the skin of DKO and control mice with a bioluminescent strain of *S. aureus* (37). *S. aureus* and *S. xylosus* are closely related Staphylococcal bacteria and both are common colonizers of the skin. Because bioluminescent *S. xylosus* is not available, examining DKO responses to *S. aureus* was used to determine if there is a generalized impairment of anti-Staphylococcal immunity in these mice. We monitored the bioluminescent signal at days 1, 3, 7, 14, and 21 after infection. In this assay, Ets1 KO mice did not show any increased susceptibility and cleared the infection with kinetics similar to wild-type mice (Figure 5D). As previously reported, mice lacking IL17RA showed a susceptibility to *S. aureus* with initially delayed clearance (Figure 5C), but were able to fully clear the infection by day 21 post-infection. On the other hand, DKO mice initially started to clear the infection and on day 3 actually showed a signal that was lower than IL17RA KO mice and comparable to that of wild-type and Ets1 KO controls. However, by day 7 the infectious signal in DKO mice increased dramatically and remained elevated until day 21 and beyond (Figure 5C). Additionally, although the initial infections were made in the skin of the upper back, in some mice the infection in DKO mice spread to the face/neck area (Figure 5D). We considered whether the susceptibility to skin infection by Staphylococcal bacteria might be due to autoimmune-mediated skin damage, given the strongly enhanced autoimmunity in DKO mice. To test whether autoimmune skin damage promotes susceptibility to Staph infection, we tested autoimmune MRL/lpr as compared to control non-autoimmune MRL/MpJ mice in the bioluminescent Staph infection model. MRL/lpr mice are known to have autoimmune-mediated skin damage, with lesions similar to human cutaneous lupus (45, 46). Despite this, we found that MRL/lpr mice could clear exogenous skin infections with *S. aureus* with the same kinetics as control MRL/MpJ mice (Figure 5E). Thus, the presence of autoimmune skin damage per se does not lead to a defect in Staph clearance.

### Alterations in γδ T cell subsets in mice lacking Ets1

In the skin of mice, several γδ T cell subsets are present, including two subsets that carry invariant TCRs, Vγ5+ dendritic epidermal T cells (DETC) and Vγ6+ dermal γδ T cells. Mice completely lacking γδ T cells have been shown to have a defect in the clearance of Staph infections from the skin, while mice lacking αβ T cells were able to clear (27). We found that DKO mice and Ets1 KO mice lack γδ TCR+ cells in epidermis (Figure 6A), while IL17RA KO and WT mice have these cells. These results were verified using flow cytometry to identify Vγ5+ and Vγ5-γδ T cells in skin. As shown in Figure 6B, DKO and Ets1 KO mice lack Vγ5+ DETC cells in skin, while IL17RA KO and WT mice have such cells. On the other hand, all genotypes of mice had Vγ5-γδ T cells in skin and these tended to be slightly increased in the skin of DKO and IL17RA KO mice.

**Figure 6.**
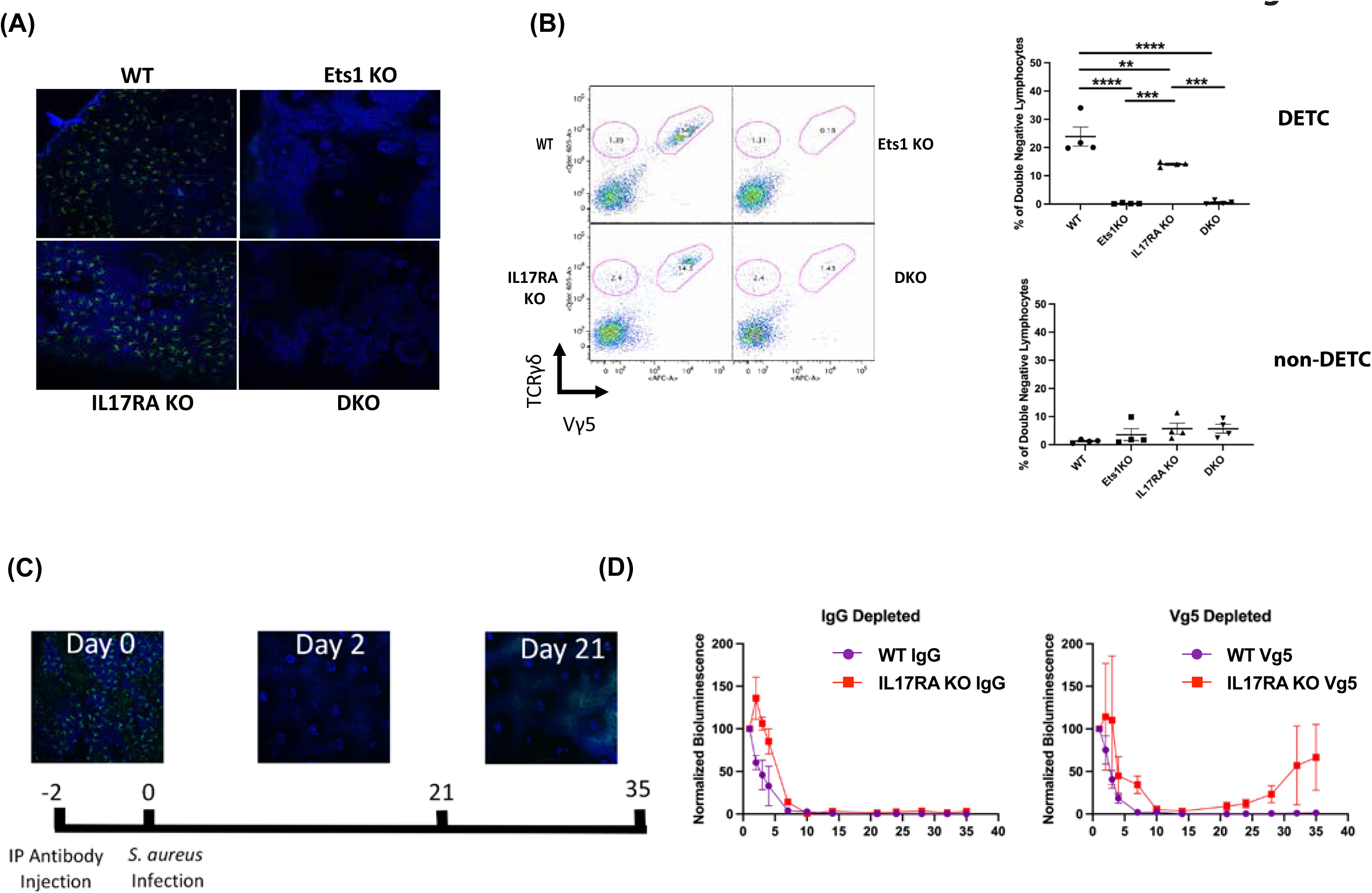
Mice lacking Ets1 lack DETC in skin and DETC are required to clear skin Staphylococcal infection. (A) Immunostaining of γδ T cells in the skin of the ear of DKO and control mice. Green staining is for the γδ TCR and blue staining is nuclei. (B) Representative flow cytometry plot for γδ T cells in the skin of mice. Plots are gated on CD4-CD8-lymphocytes and shown is staining of TCR γδ versus TCR Vγ5 (the canonical TCR of DETC). DETC are double positive for both markers and other γδ T cell subsets are positive only for TCR γδ. Graphs on the right show quantification of flow cytometry results on skin γδ T cells. N=4 mice of each genotype. (C) Immunostaining of γδ TCR+ cells (green) and nuclei (blue) before and after Vγ5+ cell depletion. Shown at the bottom is the timeline of depletion and infection. (D) Quantification of IVIS imaging of four-month-old DKO and control mice injected with either control IgG (left panel) or anti-TCR Vγ5 antibody (right panel) and then infected with bioluminescent *S. aureus*. N=2-7 mice per genotype. **p< 0.01, ***p<0.001, ****p<0.0001.

The specific role of DETC in responding to skin Staph infection is unknown. To determine if DETC might be required for clearance of Staph infection, we depleted DETC using anti-Vγ5 antibody (Figure 6C). Since DETC are the only γδ T cells that express the Vγ5 receptor, the depletion is specific to DETC and does not affect other γδ T cell subsets. DETC were depleted from both WT and IL17RA KO mice and the mice were subsequently infected with bioluminescent Staph and the infection followed with IVIS imaging. As shown in Figure 6D, IL17RA KO mice that had DETC depleted initially showed normal kinetics of clearance. However, by day 20 post-infection, the bioluminescent signal in depleted IL17RA KO mice began to increase. This was not seen in IL17RA KO mice injected with isotype control Ab or in WT mice injected with either isotype control or Vγ5 antibody. Similar to what we saw in a subset of DKO mice, we found that the infection in some IL17RA KO depleted for DETC shifted from the initial site of infection on the upper back to the face region (not shown). These results indicate that DETC cooperate with pathways induced by IL-17 signaling to mediate complete clearance of Staph infections from skin.

## Discussion

In order to explore the potential contribution of the cytokine IL-17 to the autoimmune phenotype of Ets1 KO mice, we generated double knockout mice lacking both Ets1 and the IL-17 receptor subunit IL17RA. IL17RA knockout mice are resistant to a number of autoimmune diseases including experimental autoimmune encephalomyelitis (47), collagen-induced arthritis (48), autoimmune glomerulonephritis (49) and spontaneous lupus in the BDX2 mouse model (50). This and other data support a pro-inflammatory role for signaling through IL17RA. However, a number of observations also support an anti-inflammatory role for IL-17. For instance, in autoimmune uveitis, IL-17 has been found to be protective, rather than pathogenic (51). Similarly, in sodium dextran sulfate-induced colitis, IL-17 is protective (52). In autoimmune B6.lpr mice, deletion of IL17RA leads to enhanced lymphoproliferation, though levels of anti-DNA autoantibodies are not enhanced (53). In the current study, we demonstrate that DKO mice lacking both Ets1 and IL17RA have worse autoimmune symptoms that Ets1 KO mice, suggesting that IL-17 signaling is required to limit autoimmune responses in Ets1 KO mice.

The worsened autoimmune response in DKO mice is particularly evident in the extremely enlarged skin-draining lymph nodes, which contain dramatically increased numbers and percentages of germinal center B cells, class-switched B cells, memory B cells and plasma cells. Class-switching was predominantly to the IgG1 and IgE isotypes, which is classically promoted by the cytokine IL-4. Ets1 KO mice are known to have increased numbers of T follicular helper cells that secrete IL-4 (Tfh2 cells) (22). In keeping with this, we found increased numbers of T cells with a Tfh phenotype and increased numbers of T cells that express GATA3 in spleens and lymph nodes of DKO mice. We found that there was a 2-3 fold increase in percentage of CD4+ regulatory T cells in DKO mice, whereas we did not find a change in the percentage of Tregs in Ets1 KO mice.

These data are contradictory to a previous report that found a 3-4 fold decrease in Foxp3+ Tregs in Ets1 KO mice (17). The difference between our study and the prior one may be a result of different Ets1 knockout alleles used or a difference in genetic background, age or sex of the mice analyzed. Despite the fact that there are significant increases in the numbers of Tregs in DKO mice, these cells likely have impaired function since they express lower than normal levels of Foxp3. Indeed, a previous study has shown that Tregs lacking Ets1 have reduced suppressive activity towards effector T cells (17).

In addition to the enhanced autoimmune phenotype detected in DKO mice, these mice also displayed a striking susceptibility to bacterial skin infections. DKO mice spontaneously develop skin lesions colonized by *S. xylosus*. Furthermore, four-month-old DKO mice experimentally-infected with bioluminescent *S. aureus* showed an inability to clear the infection. Given the previously-established importance of γδ T cells in controlling *S. aureus* skin infections (27), we examined γδ T cell populations in skin of DKO mice. We found that both Ets1 KO mice and DKO mice completely lack the DETC subset of γδ T cells. DETC are located within the epidermal layer and intercalate between the keratinocytes with dendritic-like cellular projections. They have been shown to have an important role in skin wound healing by production of epithelial growth factors (Fgf7, Fgf10, Igf1), inflammatory cytokines (IL-2, IFNγ) and chemokines (Ccl3, Ccl4, Ccl5, Xcl1) (54). DETC are also important for homeostatic maintenance of skin in the absence of wounding by producing IGF-1 that inhibits keratinocyte apoptosis (55). Interestingly, Ets1 KO mice have previously been shown to have a defect in skin wound healing (56). This defect was ascribed to impaired wound angiogenesis in the absence of Ets1, but may also be in part due to a lack of DETC in these mice.

Mice completely lacking all γδ T cell subsets show an impaired ability to clear skin infections with *S. aureus* (27, 57). Various subsets of γδ T cells are found in the skin, including Vγ5+ DETC and Vγ4+ and Vγ6+ cells that secrete IL-17 (58, 59). γδ T cells carrying an invariant Vγ6 chain have been shown to expand upon *S. aureus* infection, while DETC and other γδ T cell subsets are not expanded (60). That result suggests that invariant Vγ6+ cells may be important to skin immune responses to *S. aureus* infection. Vγ6+ cells make IL-17 and both IL-17 and the IL-17 receptor are required for clearance of skin *S. aureus* infections (27). Our results suggest that DETC are also one of the subsets of γδ T cells that contribute to anti-Staph immune responses, since IL17RA KO mice depleted of DETC showed enhanced susceptibility to infection. This role of DETC was only revealed in mice lacking IL17RA, but not in wild-type mice, which is consistent with the lack of observed susceptibility to *S. aureus* skin infection in Ets1 KO animals. While depletion of DETC in IL17RA mice leads to failure to clear the Staph infection, the kinetics of this response differ somewhat from DKO mice (compare figures 5C and 6D). This observation hints that there may be additional defects in the skin or in the immune response to Staphylococcal infection in DKO mice that are not mimicked by depleting DETC from IL17RA KO mice and future studies will explore this possibility.

DETC may contribute to anti-Staph immunity by promoting keratinocyte proliferation and wound healing, thus recreating the skin barrier and excluding entry of bacteria from the surface. The Vγ6+ cells that expand in response to Staph infection carry a canonical invariant TCR chain with CDR3 regions encoding the amino acid sequence CACWDSSGFHKVF (60). The invariant TCR of DETC has the same amino acid sequence in the Vγ5 CDR3 region. The TCR of DETC has been shown to recognize a self-ligand expressed by wounded or stressed keratinocytes (61). Invariant Vγ6+ cells may then respond to the same or similar TCR ligands given the identical sequence of the TCR Vγ CDR3 region. Therefore, it appears that multiple γδ T cell subsets may be involved in anti-Staph skin immunity. The DETC subset may contribute by promoting wound healing to prevent the invasion of bacteria from the skin surface, while the Vγ6+ subset may contribute by producing IL-17 that induces production of chemokines (attracting neutrophils) and anti-microbial peptides (directly killing bacteria).

While human skin lacks a direct counterpart of DETC, there are γδ T cells in human skin and they have been implicated in wound healing responses (62). These cells may contribute to the clearance of *S. aureus* skin infections by promoting wound closure. Additional data also supports a potential role for human γδ T cells in the response to *S. aureus*. In SCID mice with a humanized immune system, treatment with pamidronate, a ligand for the Vγ2Vδ2 TCR, resulted in increased clearance of intraperitoneal bacterial infections, including *S. aureus* infection (63). Recent data also show a role for human γδ T cells in promoting dendritic cell activation in the context of *S. aureus* infection and thereby resulting in increased CD4+ T cell activation (64). *S. aureus* infected APCs can also stimulate human γδ T cell production of IFN-γ (65).

A number of studies point to potential relevance of *S. aureus* skin infections in lupus patients. When the skin microbiome of lupus patients was compared to healthy controls, lupus patients showed enhanced colonization by *S. aureus* (66). In another study focusing on cutaneous lesions in lupus patients, *S. aureus* was recovered from approximately half of the lupus skin lesions, while it was not recovered from skin lesions of patients with psoriasis (67). Lupus patients whose skin lesions were colonized by *S. aureus* tended to have worse Cutaneous Lupus Disease Area and Severity Index (CLASI) scores. In addition to colonizing the skin, *S. aureus* can also be carried nasally. While lupus patients were not found to have higher rates of nasal carriage than control subjects, those lupus patients who did have nasal *S. aureus* colonization had elevated anti-dsDNA, anti-RNP, anti-SSA, and anti-SSB autoantibody titers and increased rates of kidney disease, furthering a link between *S. aureus* and lupus (68). In another study, nasal carriage of *S. aureus* was found to be associated with hypocomplementemia and with the occurrence of disease flares during the time-frame of the study (69). Together, these results implicate *S. aureus* in the induction and progression of human lupus autoimmunity. Colonization with *S. aureus* may trigger increased immune cell activation and lead to disease flares in patients with SLE. Similarly, Staphylococcal skin colonization in DKO mice may trigger enhanced immune activation and this may account for the strongly enhanced Tfh, germinal center and plasma cell responses. This is consistent with the dramatic enlargement of skin-draining lymph nodes, while interior lymph nodes and the spleen showed a less dramatic enlargement in DKO mice. In summary, the results presented in this report suggest two important conclusions: (1) that DETC function along with pathways triggered by IL-17 signaling to induce protective skin immune responses to staphylococcal skin infection and (2) that chronic skin infection by staphylococcal bacteria may trigger enhanced activation of immune cells in the draining lymph nodes leading to the intensification of an underlying autoimmune response.

## Declarations

### Ethics approval

Animal experiments were performed under the approval and guidance of the Institutional Animal Care and Use Committee (IACUC) of Roswell Park Cancer Institute protocols #UB1104M (“Ets Transcription Factors in Hematopoiesis”) and UB1300M (Skin Immunity in the Absence of Ets1 and IL17RA).

### Consent for publication

Not applicable

### Availability of data and material

Not applicable.

### Author’s contributions

MB, ACS, WL, RJ, AS, ML and LAG-S performed experiments. MB, ACS, WL, RJ, LSM, SS, EW and LAG-S designed and interpreted experiments. MB, ACS, and LAG-S wrote the manuscript. All authors read, edited and approved the final version.

## Acknowledgements

We thank Dr. Roger Plaut at the FDA for providing the bioluminescent strain of *S. aureus* for IVIS imaging. We also thank Dr. Sarah Gaffen at the University of Pittsburgh for helpful discussions on the role of IL17 in regulating immune responses to pathogens and Kirsten Smalley for help with maintaining the mouse colony.

