## Supplementary material for "Ets1 and IL17RA cooperate to regulate autoimmune responses as well as skin immunity to Staphylococcus aureus": Legends to Supplemental Figures

**Legends for Supplemental Figures**

**Supplemental Figure 1. DKO mice have increased plasma cells.** Representative flow plots showing an increase in B220-low CD138+ plasma cells in the spleen and lymph node of DKO mice. All plots are gated on total live cells.

**Supplemental Figure 2. DKO mice have increased IgG1+ B cells.** Representative flow plots showing an increase in B220+ IgG1+ class-switched B cells in the spleen and lymph nodes of DKO mice. All plots are gated on total live cells.

**Supplemental Figure 3. DKO mice have increased germinal GC B cells.** Representative flow plots showing an increase in PNA+ FAS+ germinal center B cells in the spleen and lymph nodes of DKO mice. All plots are gated on live B220+ B cells.

**Supplemental Figure 4. DKO mice have increased memory B cells.** Representative flow plots showing an increase in memory phenotype B cells (CD80+ PDL2+) in the spleen and lymph nodes of DKO mice. All plots are gated on live B220+ B cells.

**Supplemental Figure 5. DKO mice have increased T follicular helper cells.** Representative flow plots showing an increase in PD1-hi CXCR5-hi Tfh cells in the spleen and lymph nodes of DKO mice. All plots are gated on live CD4+ cells.

**Supplemental Figure 6. Both Ets1 KO and DKO mice show an increase in *in vitro* differentiation of Tfh cells.** Representative flow plots of *in vitro* differentiated Tfh cells. All plots are gated on live CD4+ cells. The upper row shows PD1 versus CXCR5 staining and the lower row shows ICOS versus BCL6 staining in the PD1-hi CXCR5-hi boxed regions shown in the upper row.

**Supplemental Figure 7. DKO mice have increased Treg cells in lymph nodes.** (A-B) Graphical quantification of the percentages of CD4+Foxp3+ regulatory T cells in spleen and lymph nodes of mice of the indicated genotypes (n=3-5 mice per group). (C) Quantification of the MFI of FoxP3 staining in the CD4+ FoxP3+ Treg population in the lymph nodes of the indicated mice. (C-D) Quantification of the numbers of CD4+GATA3+ cells and CD4+RORγt+ cells in spleen and lymph nodes of mice (n=3-5 mice per group). *$p < 0.05$, **$p<0.01$, ***$p<0.001$, ****$p<0.0001$.

**Supplemental Figure 8. Identification of *Staphylococcus xylosus* in DKO skin cultures.** (A) Pathway analysis of pathways upregulated in DKO skin as identified by RNA-sequencing. Shown are GO terms that are enriched, with the red arrow pointing to enrichment of a "*Staphylococcus aureus infection*" signature. (B) Heatmap showing strong upregulation of many cytokines and chemokines in the skin of DKO mice as compared to control mice. (C) Sequence comparison shows that the sequenced genome of the DKO Staphylococcal isolate (highlighted in green) groups with *S. xylosus* and not *S. aureus* or other Staph species. Orange highlighting represents various species of Staphylococci that are known can be the causative agents for infections in humans.
