## Supplemental Figures for "Ets1 and IL17RA cooperate to regulate autoimmune responses as well as skin immunity to Staphylococcus aureus"

### Supplemental Figure 1

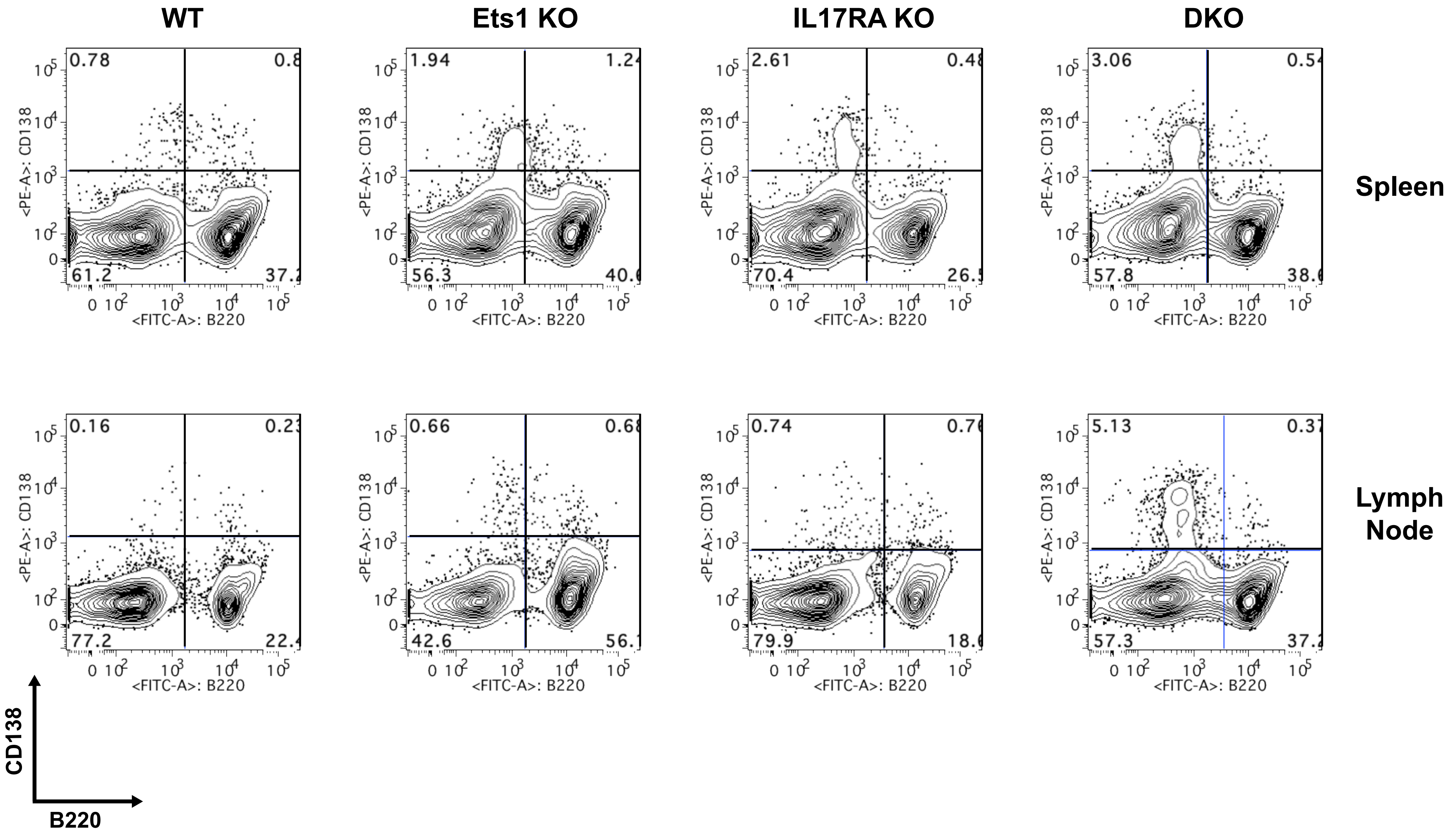

**Supplemental Figure 2**

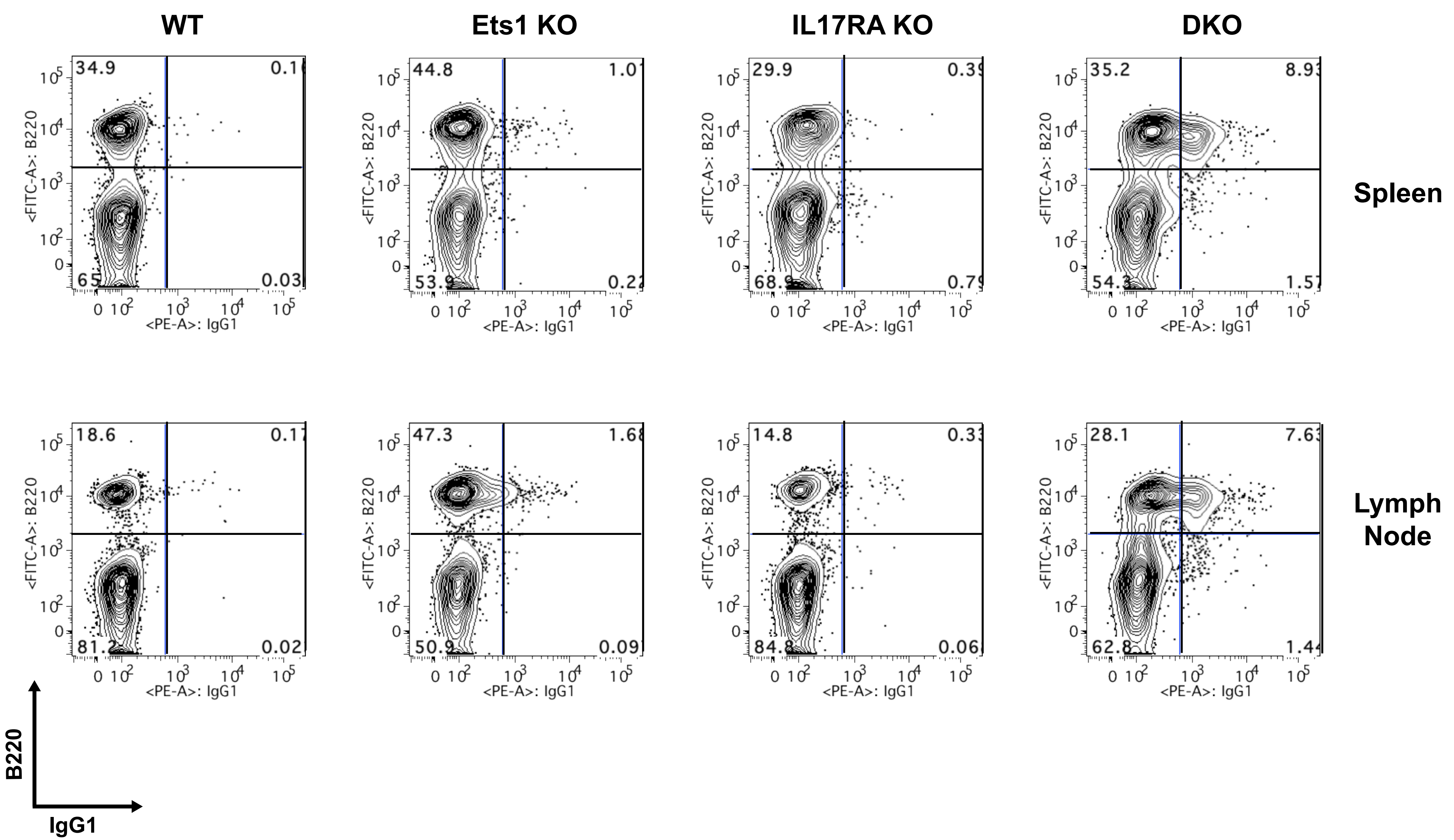

### Supplemental Figure 3

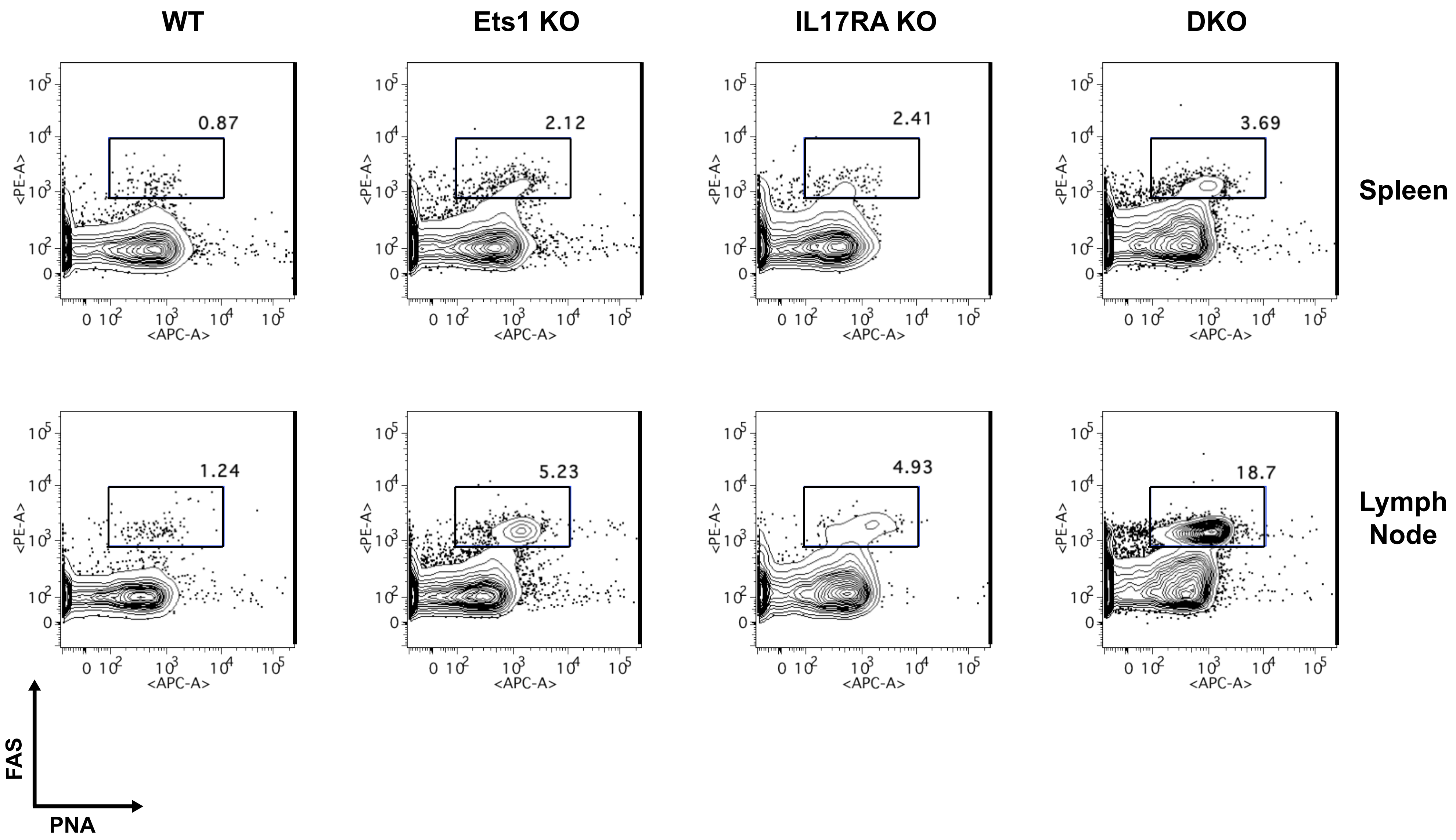

### Supplemental Figure 4

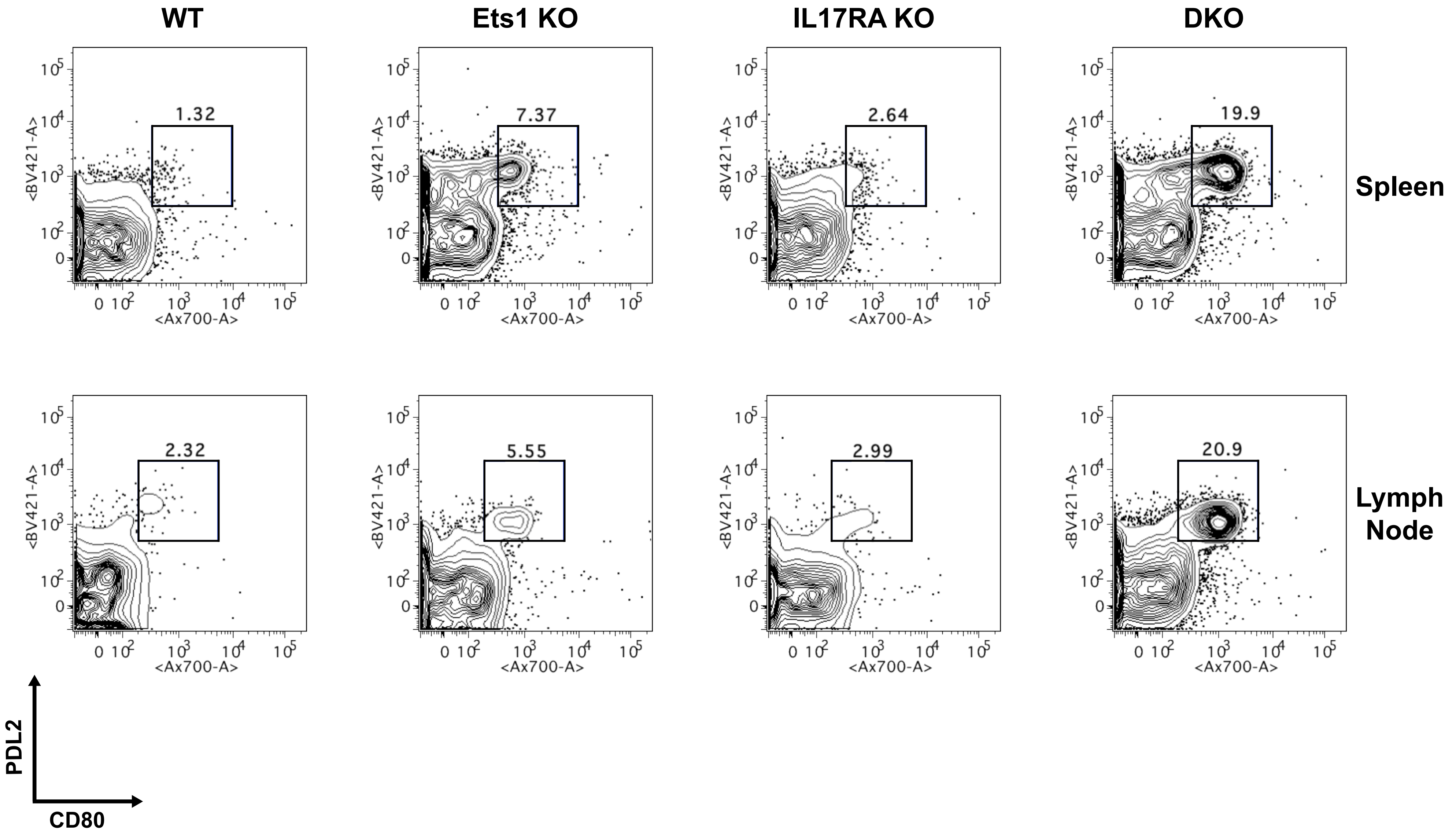

#### Supplemental Figure 5

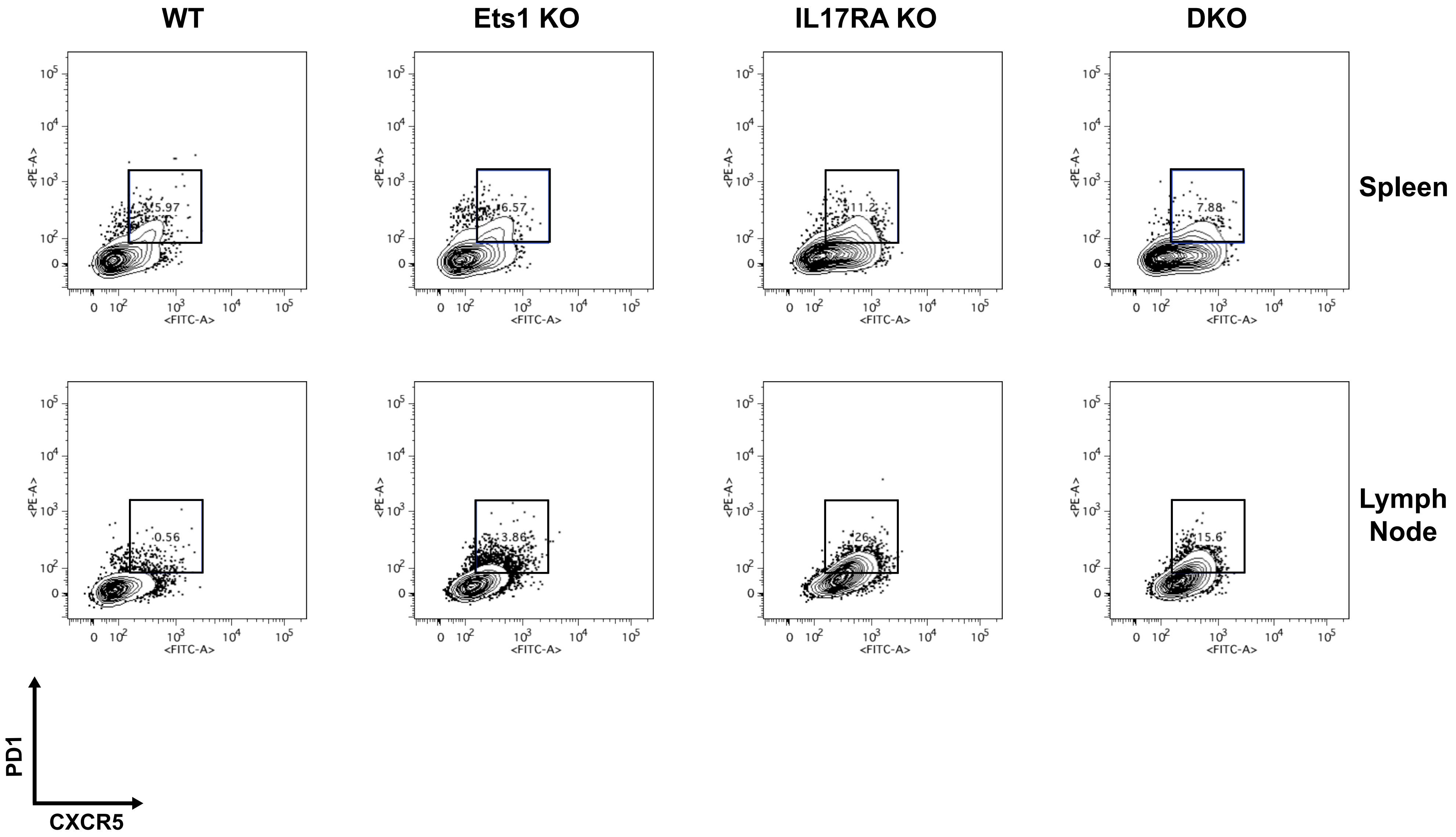

#### Supplemental Figure 6

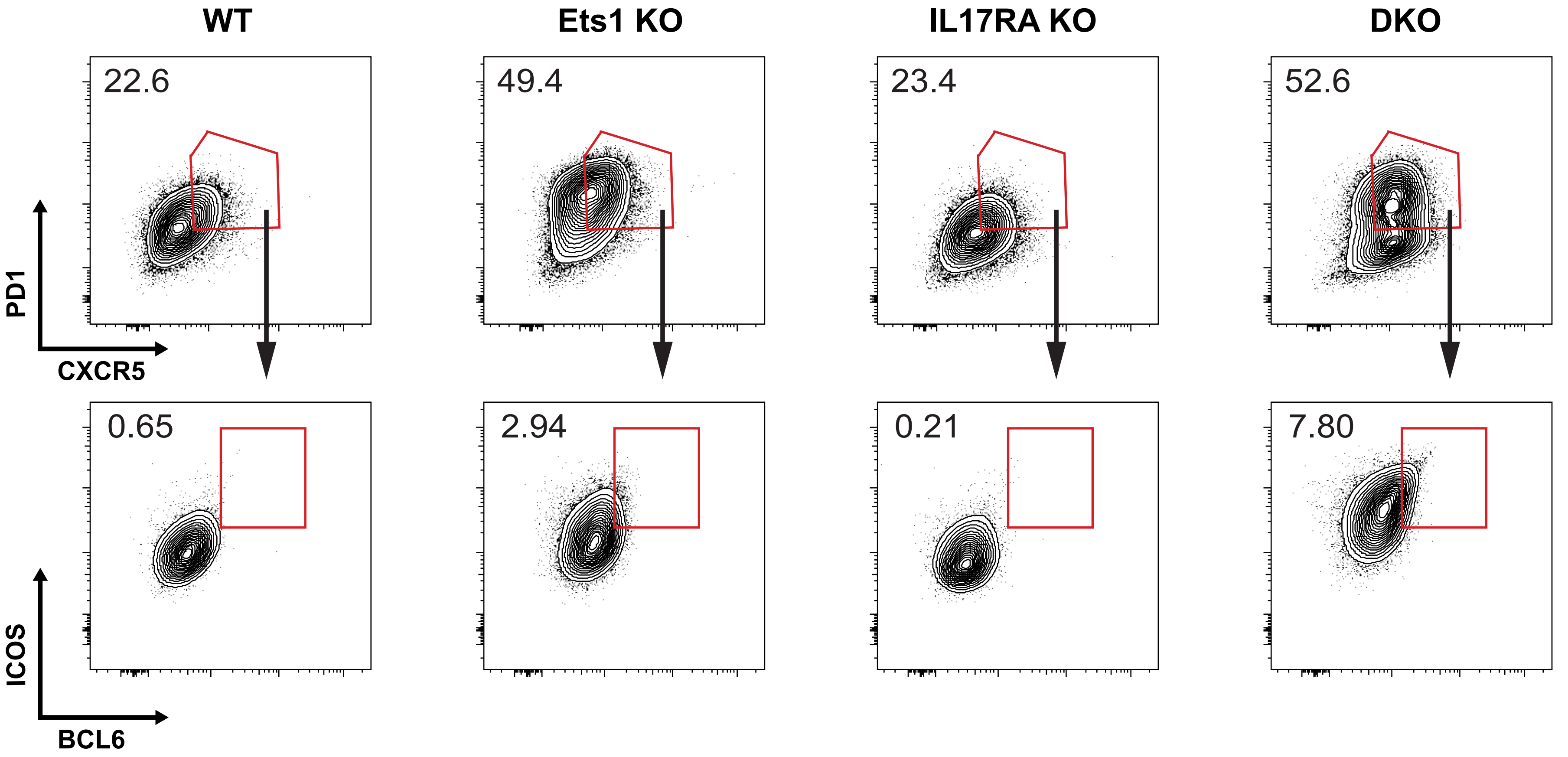

#### Supplemental Figure 7

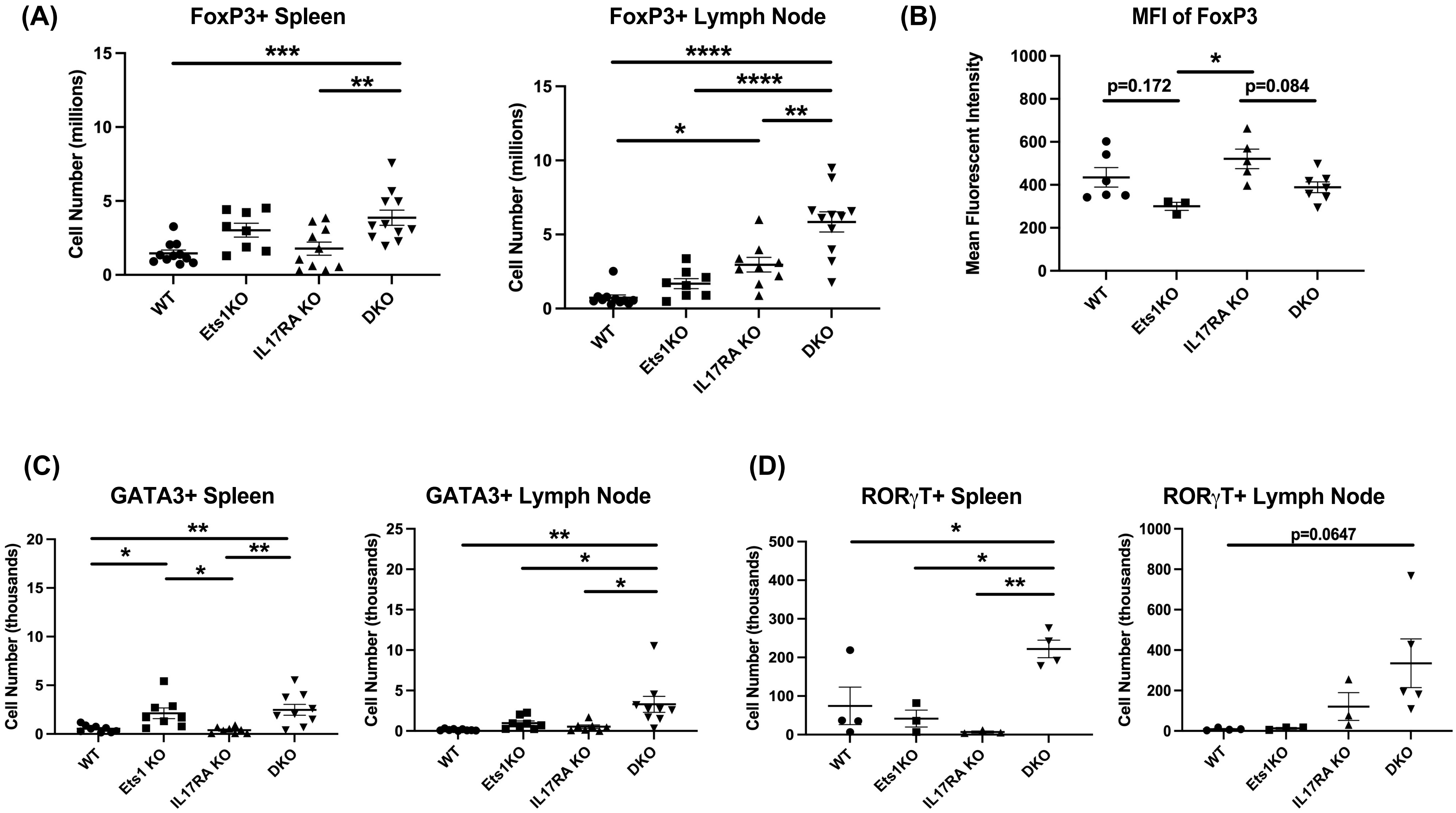

#### Supplemental Figure 8

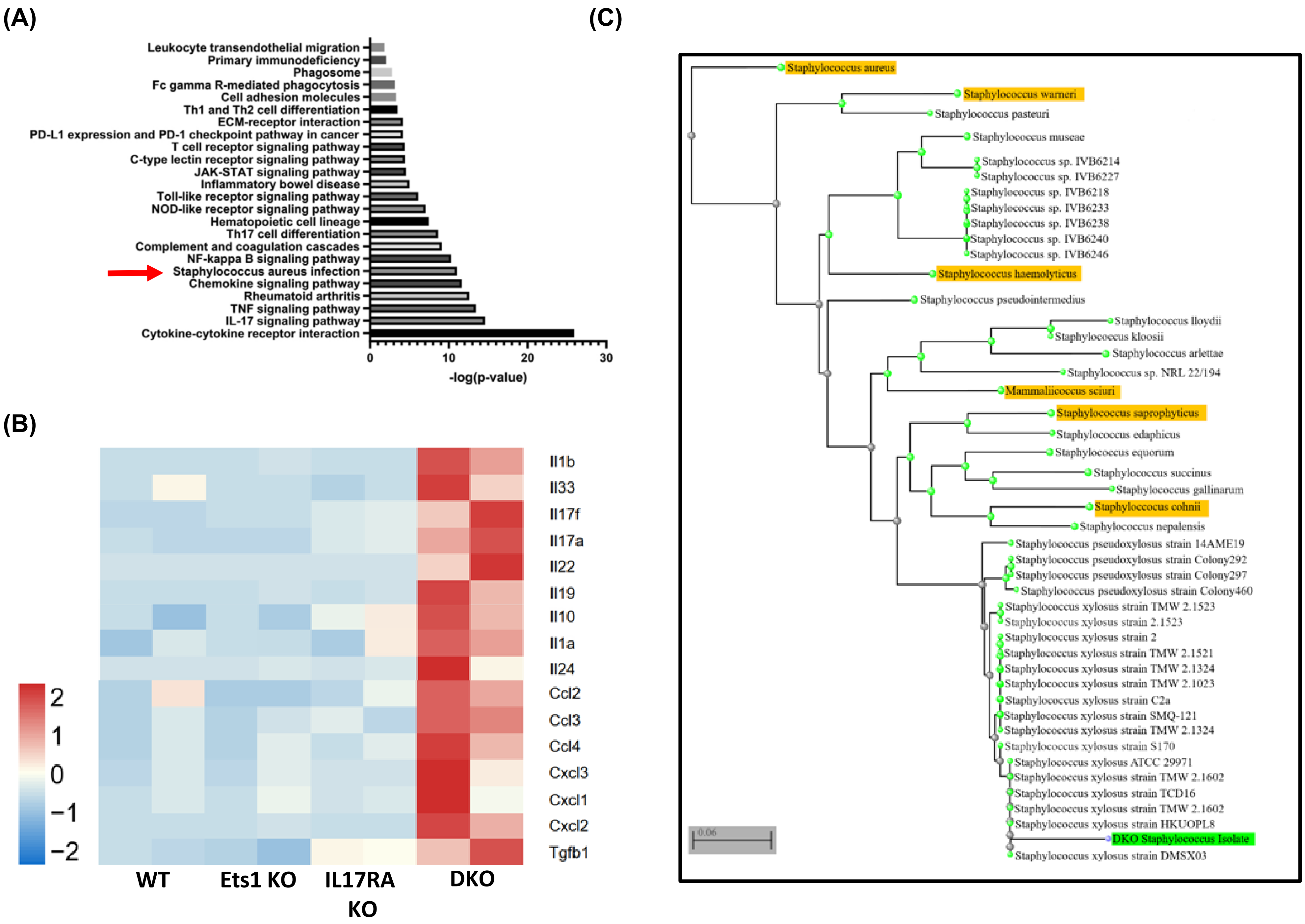
